# How gut hormones shape reward: a systematic review of the role of ghrelin and GLP-1 in human fMRI

**DOI:** 10.1101/2022.11.30.518539

**Authors:** Corinna Schulz, Cecilia Vezzani, Nils B. Kroemer

## Abstract

The gastrointestinal hormones ghrelin and glucagon-like peptide-1 (GLP-1) have opposite secretion patterns, as well as opposite effects on metabolism and food intake. Beyond their role in energy homeostasis, gastrointestinal hormones have also been suggested to modulate the reward system. However, the potential of ghrelin and GLP-1 to modulate reward responses in humans has not been systematically reviewed before. To evaluate the convergence of published results, we first conduct a multi-level kernel density meta-analysis of studies reporting a positive association of ghrelin (*N_comb_*= 353, 18 contrasts) and a negative association of GLP-1 (*N_comb_* = 258, 12 contrasts) and reward responses measured using task functional magnetic resonance imaging (fMRI). Second, we complement the meta-analysis using a systematic literature review, focusing on distinct reward phases and applications in clinical populations that may account for variability across studies. In line with preclinical research, we find that ghrelin increases reward responses across studies in key nodes of the motivational circuit, such as the nucleus accumbens, pallidum, putamen, substantia nigra, ventral tegmental area, and the dorsal mid insula. In contrast, for GLP-1, we did not find sufficient convergence in support of reduced reward responses. Instead, our systematic review identifies potential differences of GLP-1 on anticipatory versus consummatory reward responses. Based on a systematic synthesis of available findings, we conclude that there is considerable support for the neuromodulatory potential of gut-based circulating peptides on reward responses. To unlock their potential for clinical applications, future studies may move beyond anticipated rewards to cover other reward facets.

**Graphical Abstract:** 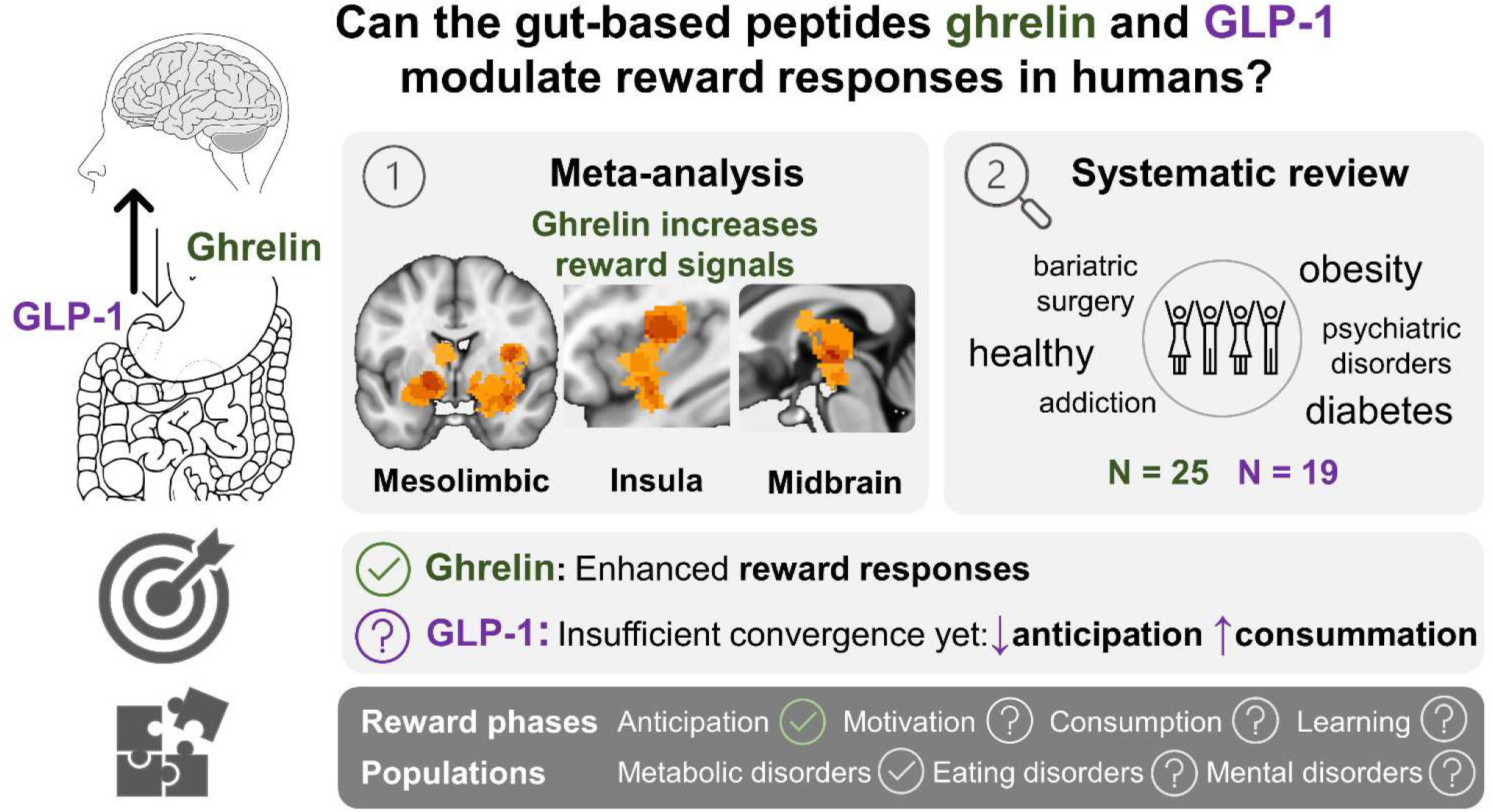

## Introduction

Gastrointestinal hormones, such as ghrelin and glucagon-like peptide-1 (GLP-1), control various functions within the digestive system while communicating with our central nervous system (CNS) as part of the gut–brain axis. The gut–brain axis enables bidirectional communication between the brain and the gastrointestinal system, involving neuroendocrine pathways, the vagus nerve, immune factors, and microbiota (Carabotti et al., 2015; Cryan et al., 2019; Holzer & Farzi, 2014; Mayer et al., 2022). Whereas peripheral factors from the gastrointestinal system are known to exert influence on “homeostatic” feeding areas, such as the brainstem–hypothalamic circuit, there is evidence that they can also—directly or indirectly—influence the brain’s reward system (Klockars et al., 2021; Liu & Borgland, 2015). The reward system is a distributed network with its key structures including the ventral striatum, the anterior cingulate cortex, the orbitofrontal cortex, the ventral pallidum, and the midbrain dopamine neurons (Haber & Knutson, 2010). However, reward is not a monolithic entity (Berridge & Robinson, 2003). Instead, reward responses reflect distinct psychobiological components such as motivation (“wanting”), hedonics (“liking”), and learning (Berridge et al., 2009). Distinguishing these facets of reward-related behavior is crucial for clinical applications as symptoms and their treatment may map onto specific facets (Husain & Roiser, 2018; Treadway & Zald, 2011). Accordingly, preclinical research has established crucial mechanisms of neurotransmission within the reward system that differentiate between motivational aspects of reward (i.e., dopamine) rather than the pleasure (i.e., opioids and endocannabinoids) itself (Berridge & Kringelbach, 2015; Volkow et al., 2017). Notably, several regions in the reward system co-express receptors for neurotransmitters and neuropeptides, allowing metabolic state to directly influence neural signaling (e.g., dopamine, acetylcholine; Ferrario et al., 2016; Liu & Borgland, 2015). To conclude, the role of gastrointestinal hormones extends beyond local functions within the digestive system and to communication with the CNS, potentially shaping reward-related processes according to metabolic demand.

In this systematic review and meta-analysis, we address which role ghrelin and GLP-1 play in modulating reward responses as measured with task fMRI in humans. Because of their opposite functions on food intake and metabolic targets (Engelstoft & Schwartz, 2016), both hormones have gained traction as pharmacological targets for treatment of metabolic (i.e., obesity and type 2 diabetes (T2D)) and eating disorders (Atalayer et al., 2013; Drucker, 2022; Gupta et al., 2021; McElroy et al., 2018). A recent review concludes that ghrelin and GLP-1 also show opposing effects on food reward-associated behaviors in rodent and humans (Decarie-Spain & Kanoski, 2021). Here, we conducted the first systematic review and meta-analysis of human fMRI studies to investigate whether the reward system is robustly modulated by ghrelin and GLP-1. Furthermore, in our systematic review, we emphasize important differences in findings between different clinical populations and different reward phases that may help inform future studies as well as clinical applications. We conclude our review on the modulatory role of ghrelin and GLP-1 in reward processing by highlighting gut-based interventions that can be used to treat motivational and emotional dysfunctions across a range of mental and metabolic disorders.

### Ghrelin and its central effects

Ghrelin is a 28-amino-acid orexigenic peptide hormone primarily produced and secreted by entero-endocrine cells of the stomach (Kojima et al., 1999). Ghrelin’s functions include regulating energy homeostasis, controlling central homeostatic and hedonic feeding, preventing hypoglycemia under famine, and protecting from metabolic stress and metaflammation (Yanagi et al., 2018). Since circulating ghrelin levels rise upon fasting and drop after meal intake (Cummings et al., 2001), it was initially described as the ‘hunger hormone’. Ghrelin stimulates food intake by acting on hypothalamic neuropeptide Y (NPY)/ agouti/related peptide (AgRP) and proopiomelanocortin (POMC) neurons (Chen et al., 2004).

How circulating ghrelin gains access to the CNS is dependent on its unique posttranslational process in which the serine-3 is *n*-octanoylated (Yanagi et al., 2018). This modification is essential for ghrelin’s biological activity since acylated ghrelin has a much higher affinity for the growth-hormone secretagogue type 1a receptor (GHS-R1a), the primary target in the brain. GHS-R1a mRNA is expressed in the periphery (e.g. pancreatic islets, adrenal gland, thyroid, lung, liver, kidney, intestine, adipose tissue), as well as in the CNS (e.g. hypothalamus, hippocampus, substantia nigra, ventral tegmental area (VTA), nucleus of the solitary tract (NTS)) (Cornejo et al., 2018; Yanagi et al., 2018). Whereas the role of des-acyl (des-octanoyl) ghrelin in activating GHS-R1a is still debated, it has been suggested that des-acyl ghrelin can influence food intake by mechanisms independent of GHS-R1a (Toshinai et al., 2006) and by modulation of insulin sensitivity (Heppner et al., 2013). Since there is limited evidence for central ghrelin synthesis, circulating ghrelin needs to be transported into the brain for central action on GHS-R1a. This can be achieved by crossing the blood-brain-barrier involving saturable mechanisms (Banks et al., 2002), by diffusing through fenestrated capillaries at circumventricular organs (Uriarte et al., 2021), or by crossing the blood-cerebrospinal barrier (Uriarte et al., 2019). Work in animals provides evidence for each of these putative pathways, supporting that ghrelin can act locally in the hypothalamus (i.e., paraventricular nucleus, arcuate nucleus), NTS, and potentially, in the hippocampus and VTA (Perello et al., 2019). Yet, conflicting evidence challenges these putative mechanisms, calling for future work on how ghrelin modulates function in the human brain (Perello et al., 2019). In addition to direct signaling, ghrelin has also been shown to affect food intake and metabolism at least partly via vagal afferent neurons in the nodose ganglion that express GHRS (Davis et al., 2020; Le Roux et al., 2005). To summarize, there are several complementary pathways for ghrelin to influence central signaling, which may facilitate tuning of the reward system.

Ghrelin’s potential to increase reward responses was discovered in rodent studies investigating motivated behaviour as well as dopaminergic transmission. For example, both central and peripheral administration of ghrelin increased the willingness of (male) mice to work for obtaining food rewards in an operant conditioning paradigm (Skibicka et al., 2012). Whereas this study did not directly assess dopamine, they reported increased dopamine receptor gene expression in the VTA. Other studies in (male) mice have shown the effects of ghrelin on dopamine transmission via a variety of administration routes. Specifically, local ghrelin injections to the VTA and laterodorsal tegmental nucleus as well as central administration to the lateral or third ventricle increased extracellular dopamine in the nucleus accumbens (NAc; Cone et al., 2014; Jerlhag et al., 2006; Kawahara et al., 2009). Also, peripherally administered ghrelin has been shown to augment extracellular dopamine in the NAc (Jerlhag, 2008; Quarta et al., 2009) and in the VTA during sexual interactions (Prieto-Garcia et al., 2015). To conclude, work in rodents provides conclusive support for the idea that ghrelin modulates motivated behavior via increased dopaminergic signaling in the reward system.

### GLP-1 and its central effects

GLP-1 is an incretin hormone which is produced in L-cells in the intestine, α-cells in the pancreatic islet, and in neurons in the NTS (D’Alessio, 2016). GLP-1’s peripheral effects include insulin secretion, gut emptying, and inhibition of glucagon secretion. Opposite to ghrelin, GLP-1 is secreted in response to food intake and decreases meal size (Krieger, 2020). Consequently, GLP-1 analogs are used for treatment of T2D and the GLP-1 analogs liraglutide and, more recently, semaglutide have been approved by the Food and Drug Administration to support weight loss in obesity (Trapp & Brierley, 2022). Like GHRS-R1a, the G protein-coupled GLP-1 receptor (GLP-1R) is also expressed in various regions of the CNS, including the hypothalamus, thalamus, brainstem, and cortical regions such as the insula, opercular cortex, frontal pole, and paracingulate gyrus (Drucker, 2022; https://neurosynth.org/genes/GLP1R/), thereby allowing GLP-1 to affect central signaling.

Similar to ghrelin, GLP-1 exerts central effects via complementary mechanisms acting on GLP-1R (Kanoski et al., 2016). Circulating GLP-1 can cross the blood-brain barrier and the success of GLP analogs in reducing food intake and body weight partly involve penetration of the blood-brain barrier (Kastin et al., 2002). Interestingly, GLP-1 is also produced directly in the CNS by preproglucagon neurons, predominantly in the brainstem (D’Alessio, 2016; McLean et al., 2021). While GLP-1 can bind to receptors on circumventricular regions (e.g., area postrema), it is yet unclear to what extent gut-derived or brain-derived GLP-1 contribute to central GLP-1R activation (Trapp & Brierley, 2022). Another way of action for gut-derived GLP-1 includes indirect signaling via the vagus nerve. Specifically, GLP-1Rs on vagal afferent neurons have been shown to contribute to the incretin effects of GLP-1 shortly after a meal, gastric emptying, and glycemia (Krieger et al., 2015). Altogether, circulating GLP-1 can affect central signaling by complementary pathways, thereby supporting satiety-related signaling.

In contrast to ghrelin, GLP-1 has been shown to decrease reward responses in rodents. A GLP-1 agonist reduced conditioned place preference for food rewards as well as motivated behavior for sucrose in a progressive ratio operant-conditioning paradigm in rodents via two key nodes of the reward system: the VTA and NAc (Dickson et al., 2012). More direct evidence for GLP-1 modulating dopamine transmission comes from a study showing that a GLP-1 agonist also reduced phasic dopamine response in the VTA in rats (Konanur et al., 2020). In addition to decreasing food intake, GLP-1 has also been shown to decrease drug intake, self-administration, conditioned place preference, and seeking in animal models (Eren-Yazicioglu et al., 2021), suggesting that GLP-1 can decrease reward responses for food rewards and beyond.

As established in animals, ghrelin and GLP-1 are well positioned as signaling agents within the gut–brain axis to regulate food intake and modulate motivational and reward-related behavior by increasing and decreasing reward responses, respectively. Although there is evidence that ghrelin and GLP-1 modulate reward responses in humans as well (Decarie-Spain & Kanoski, 2021), this has not been systematically reviewed before. We hypothesized to find consistent changes in regions of the reward system that are modulated by ghrelin and GLP-1 across human fMRI studies. Specifically, we hypothesized to find

1. increased reward responses after administration of ghrelin and/or positive correlations with plasma ghrelin levels, and
2. decreased reward responses after administration of GLP-1 agonists and/or negative correlations with plasma GLP-1 levels.

## Methods

### Search

The selection process consisted of two searches for English-language articles with no year limit. The first search was conducted on Web of Science with the following key terms: “*ghrelin OR glp-1 OR glucagone-like*) AND reward* AND (fmri OR mri OR neuroimag\**” on August 10^th^, 2022. This yielded 82 results. After screening of titles and abstract, 46 papers were selected as relevant for the review. A second search was conducted on PubMed on August 23^rd^, 2022, with the following terms: “*((ghrelin) OR (glp-1)) OR (glp1)) OR (glucagone-like)) AND (reward) AND ((fmri) OR (mri) OR (neuroimaging))*”, generating 76 results. After screening of titles and abstract, and removal of the results overlapping with the previous search, 16 additional papers were selected as relevant for the review.

### Selection Criteria: systematic review

Inclusion criteria for the systematic review were the collection of neuroimaging (i.e., fMRI) data during exposure or reaction to reward-related stimuli in humans, combined with the measurement of endogenous levels of ghrelin or GLP-1 and/or administration of an exogenous GHS-R or GLP-1R agonist or antagonist. Papers were excluded if they were review papers, if the study was conducted only on animals, if there was no measurement of ghrelin or GLP-1, or administration of an agonist or antagonist for the respective receptors, or when the hormonal measure or manipulation was not related to the reward response. Figure 1 provides an overview of the article selection process.

**Figure 1:**
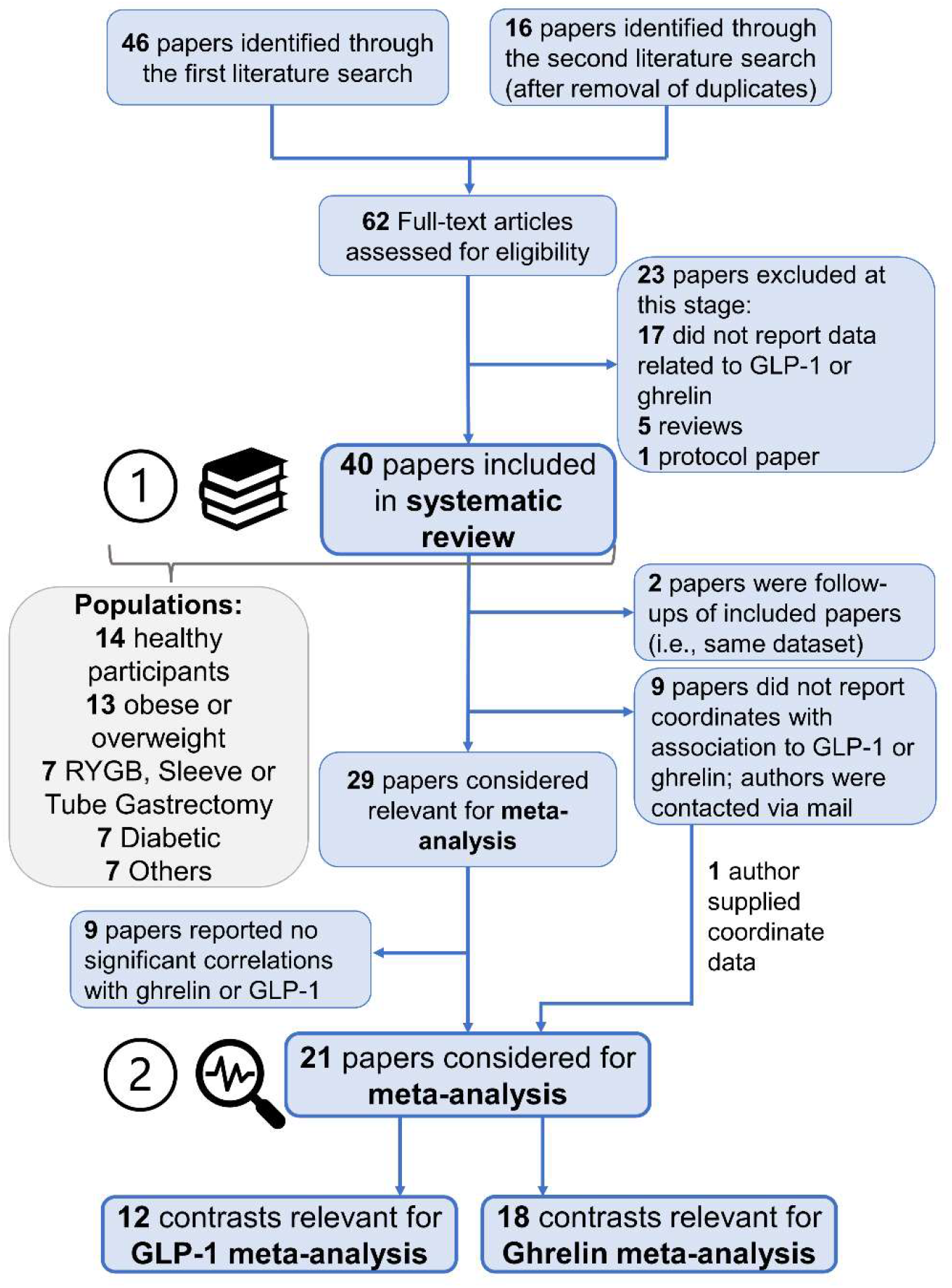
PRISMA flow diagram of the article selection process for the (1) systematic literature review and the (2) meta-analysis.

22 papers were excluded from the initial database search, leaving a total of 40 papers for the review. Two more papers were excluded from the review since because they reported data from the same sample as another included paper. The reviewed papers focused on different populations: the most represented populations where those of healthy participants (14 studies) and individuals with overweight and obesity (13 studies). Seven studies investigated patients undergoing bariatric surgery, Roux-en-Y Gastric bypass, and sleeve or tube gastrectomy, while seven others focused on patients with T2D. Finally, the remaining papers studied patients with alcohol use disorder (2), anorexia nervosa (1), acquired hypothalamic damage (1), Prader-Willi syndrome (1), and major depressive disorder (1).

An overview table of included studies is available in the Supplementary Material (S1) and online with options to filter the database (https://tinyurl.com/gut-brain-reward). The table includes information on first author and year of publication (1), paper title (2), whether the study was included in the meta-analysis (3) or systematic review (4), a tag indicating whether ghrelin or GLP-1 was investigated (5), specification of the hormonal assessment and/or manipulation (i.e., whether plasma acyl-, desacyl-, total ghrelin, GLP-1 was measured, which substance was administered (if any)) (6), type of reward phase assessed during fMRI (7), type of reward (e.g., food, money, alcohol) (8), and sample population(s) (9).

### Inclusion criteria: meta-analysis

For the meta-analysis, we included all studies that reported peak coordinates for any of the following contrasts in association with either ghrelin or GLP-1: food > non-food, high-caloric > low-caloric food, highly palatable food > low palatable food. If multiple contrasts were available, we chose food > non-food. Associations included either correlations of blood plasma hormone levels with the task contrasts or interventions where hormone levels were directly changed (i.e., injection of ghrelin, GLP-1 agonists, or GLP-1 antagonists). Relevant coordinates were reported in 14 papers focusing on ghrelin and 7 papers focusing on GLP-1. Emails were sent to the remaining authors of the identified papers, resulting in one paper being added to the ghrelin analysis. Eight papers did not report any significant correlation with ghrelin or GLP-1, while two papers were excluded from the meta-analysis because they reported follow-up analyses of data that had already been included.

## Statistical Analysis

### Meta-analysis

Meta-analysis was performed in Matlab R2021a using the multi-level kernel density meta-analysis (MKDA) toolbox from CANLab (Wager et al., 2007; 2009; Wellcome Department of Cognitive Neurology, 2005) that uses tools from Statistical Parametric Mapping (https://www.fil.ion.ucl.ac.uk/spm/). MKDA is an improved version of the kernel density analysis, which uses the proportion of studies that activate in a region (rather than the number of peaks) as the test statistic (Wager et al., 2007). Multilevel here indicates that peaks are nested within studies. The advantage is that MKDA allows to generalize to new studies as it tests for the null hypothesis that activations across studies are not spatially consistent.

For the meta-analysis, different worksheets were created based on the assessed hormone (ghrelin or GLP-1) and effect direction (positive or negative) (Supplementary Material S2). These sheets contain:

1. MNI/Talairach peak coordinates, with separate columns for x, y and z
2. First author and year of publication
3. A unique integer for each contrast map
4. Sample size
5. Indication of whether fixed or random effects were used for the analysis
6. coordinate system (either MNI or Talairach)

To estimate the effects of gut-derived peptides on brain responses, we ran separate analyses for ghrelin and GLP-1 effects on brain responses during reward tasks. We selected peak coordinates for the contrasts between food and non-food outcomes, or contrasts for high-caloric food and low-caloric food or highly palatable food and low palatable food whenever the food vs. non-food contrast was not reported. The coordinates were extracted along with the effect direction (positive association with ghrelin, negative association with GLP-1). The choice to report coordinates relative to a contrast map rather than a study was made to maximize independent contrast maps (e.g., if studies reported contrasts for different reward phases). Significant activation was therefore identified based on the number of contrast maps activated near a voxel rather than the direct number of peaks reported around that voxel (Wager et al., 2009). Coordinates for peak effects (Figure 2, Panel a) from each contrast map were convolved with a spherical kernel (*r* = 10mm) and thresholded at 1 (indicator maps). Averaging the indicator maps, weighted by sample size, yielded a density of effects map across studies (Figure 2, Panel b; Kaiser et al., 2015). The density map shows the proportion of studies in which a positive or negative association of ghrelin or GLP-1 with reward responses was observed within 10 mm of each voxel.

**Figure 2.**
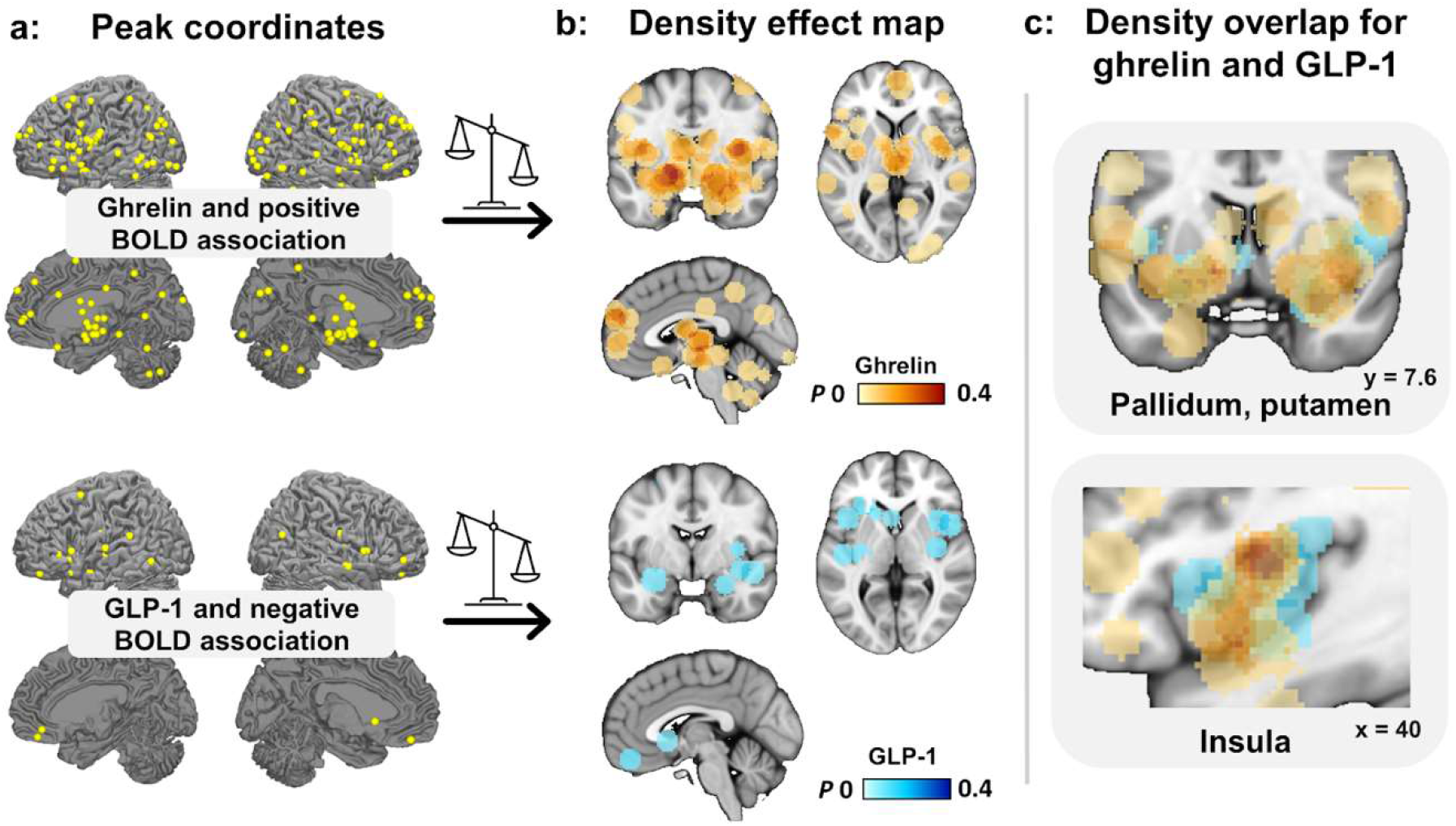
Extracted peak coordinates, density effect maps, and overlap for studies showing a positive association with ghrelin and a negative association with GLP-1. Multi-level kernel density meta-analysis was separated for coordinates reporting a positive association of ghrelin and BOLD activation (red) and studies reporting a negative association of GLP-1 and BOLD activation (blue). **a**: Peak coordinates were extracted from relevant papers, which were then convolved with a spherical kernel, and thresholded at a maximum value of 1. **b**: Convolved peak coordinates across studies are weighted by sample size to yield density effect maps. **c**: The overlap between density maps for ghrelin and GLP-1 studies is shown for the insula and striatal regions. Please note that density effect maps are uncorrected for multiple comparisons.

**Figure 3.**
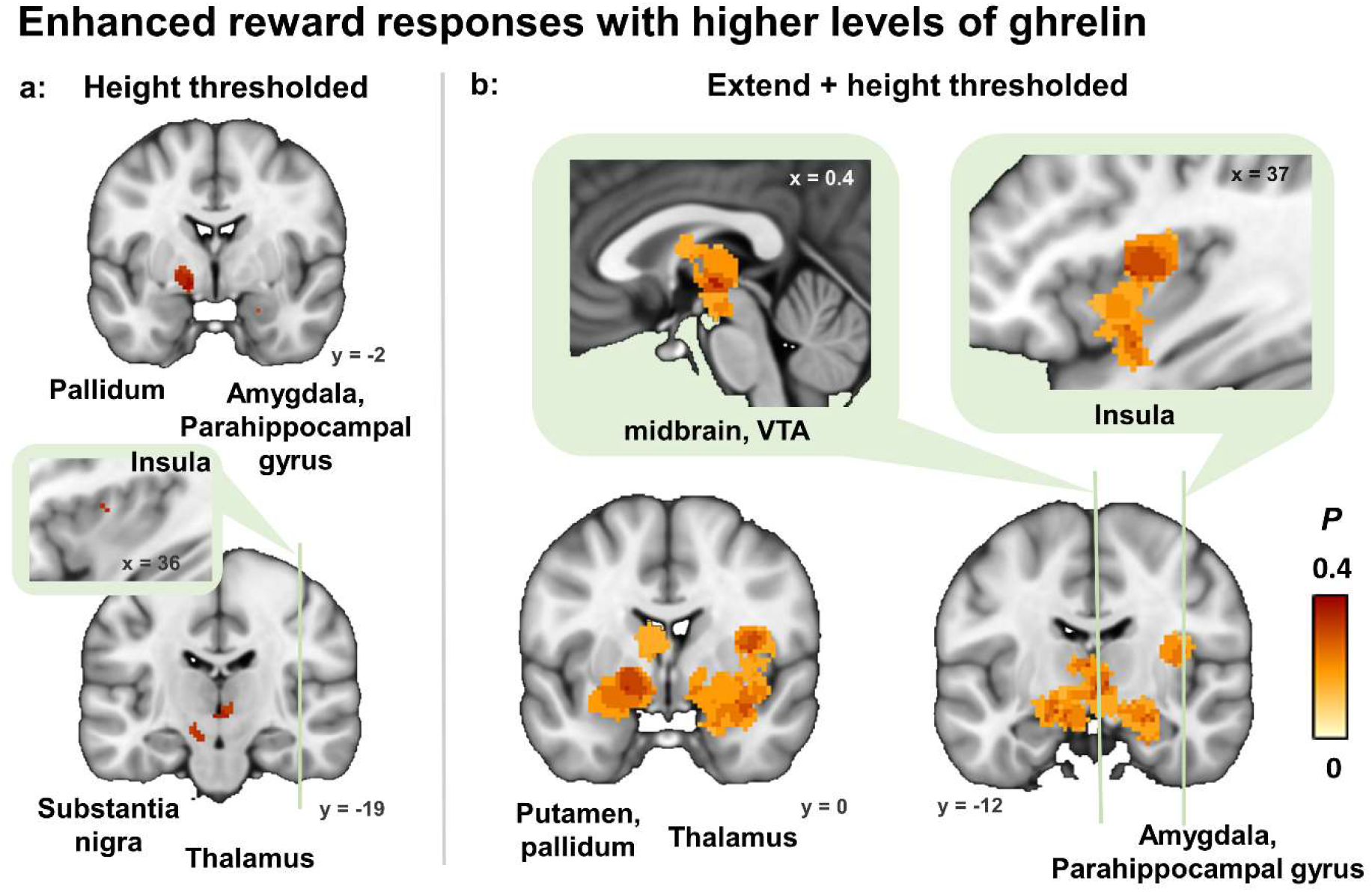
Ghrelin robustly modulates reward response increases across studies. Regions that survive correction in a multi-level kernel density meta-analysis of studies reporting a positive association of ghrelin and BOLD activation are shown. **a**: Clusters that surpass a FWE height-correction (*p* < .05). **b**: Clusters that surpass a FWE extent-based correction (*p* < .01) are shown in addition to the height-corrected clusters. *Maximum P* is the test statistic, indicating the maximum proportion of studies exhibiting the effect at the peak density weighted by sample size. VTA = ventral tegmental area.

Finally, the familywise error rate (FWE) threshold to correct for multiple comparisons was calculated through iterations using a Monte Carlo permutation procedure. Here, the significant effects from indicator maps for each statistical contrast map were compared with a null hypothesis (i.e., *activations across studies are not spatially consistent*) over 10,000 iterations (Wager et al., 2009). We report thresholded maps based on a FWE extent-corrected threshold of *P* <.01 (density at neighbouring voxels exceeds maximum expected in that cluster size by change) and include information on the FWE height-corrected threshold of *P* <.05 (density at single voxel exceeds the maximum expected across the entire brain by chance) as complementary information (Kaiser et al., 2015).

## Results

The meta-analysis and systematic review were conducted following the Preferred Reporting Items for Systematic Reviews and Meta-Analyses (PRISMA) guidelines (Page et al., 2021). After further examination of the included papers, the literature searches yielded a final result of 29 studies considered as relevant for the systematic review and 21 studies suitable for the meta-analyses. From these 21 studies, we extracted a total of 36 contrasts and 265 peak coordinates. We separated meta-analyses for ghrelin and GLP-1, as well as for coordinates reporting a positive or a negative association of ghrelin and GLP-1 with reward responses, since coordinate-based meta-analysis test for spatial convergence (Müller et al., 2018).

### 1. Meta-analysis

#### 1.1. Density maps for ghrelin and GLP-1 overlap in reward regions

For ghrelin, peak coordinates from 18 different contrasts had a positive association with reward-related brain responses (Figure 2, Panel a), whereas coordinates from only 7 different contrasts had a negative association. With respect to GLP-1, coordinates from 12 different contrasts had a negative association with reward-related brain responses (Figure 2, Panel a) and 3 contrasts had a positive association. Since the number of available contrasts for the non-hypothesized direction is too low, we will focus in the meta-analysis on the hypothesized direction for ghrelin and GLP-1.

The 18 contrasts that showed a positive association for ghrelin stemmed from 14 different studies, yielded 184 peak coordinates from a combined sample size of 353 individuals. The 12 contrasts that showed a negative association for GLP-1 stemmed from 7 different studies, yielded 26 peak coordinates and a combined sample size of 258 individuals. Across the brain, overlap between the resulting density maps for ghrelin and GLP-1 was low (Jaccard and Dice coefficients <.2). However, ghrelin and GLP-1 density maps overlapped in key regions of the reward circuit such as the globus pallidus, putamen, NAc, caudate, dorsal mid insula (Figure 2, Panel c).

#### 1.2. Ghrelin MKDA: elevated reward responses with high ghrelin

To estimate the hypothesized reward-enhancing effect of ghrelin, we conducted a meta-analysis of positive associations with ghrelin. This yielded seven major clusters of activation that surpassed the FWE extent-corrected threshold (*p* < .01; Table 1, Figure 2, Panel b). The largest cluster included the globus pallidus, right dorsal mid insula, claustrum, putamen, hypothalamus, left NAc, and thalamus. The cluster showed a maximum *P* (i.e., the maximum proportion of studies exhibiting the effect at the peak density weighted by sample size) of 0.38 at the left globus pallidus. Two smaller clusters were identified around the thalamus with a maximum *P* of 0.17. Additional clusters were identified around the putamen, the right parahippocampal region and amygdala, the red nucleus in the midbrain, and the substantia nigra. Clusters that surpassed the height-corrected threshold (*p* < .05) were smaller, but encompassed similar regions (Figure 2, Panel a; Supplementary Material S3). All maps (i.e., density effect map and thresholded maps) can be inspected and downloaded at https://identifiers.org/neurovault.collection:13247.

**Table 1.**
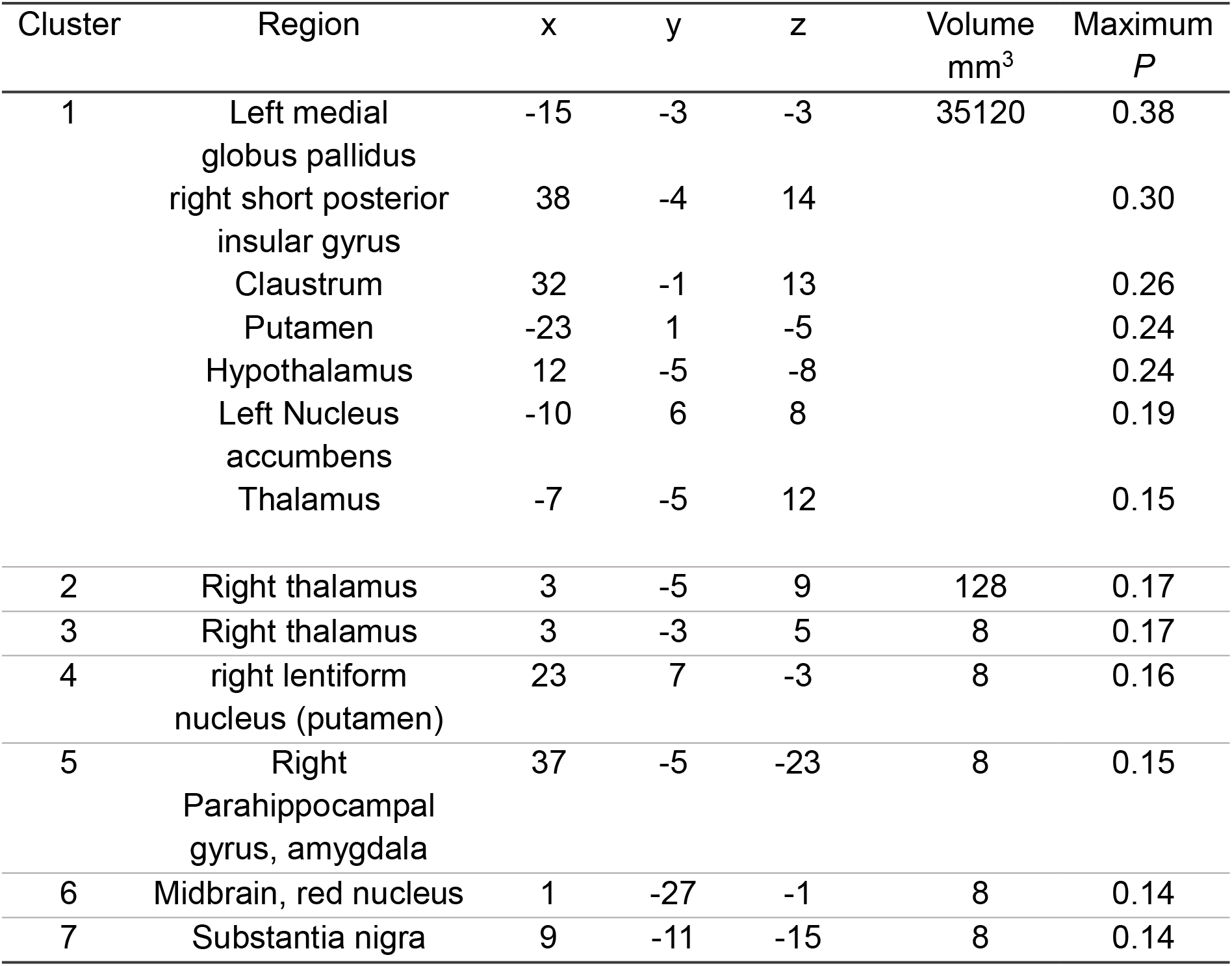
Results of the multi-level kernel density meta-analysis reporting a positive association of ghrelin and BOLD activation. Clusters that show a positive association between ghrelin and reward responses, which surpass the medium extent-based threshold (*p* < .01). Coordinates are Montreal Neurological Institute standard stereotaxic spaces. *Maximum P* is the test statistic indicating the maximum proportion of studies exhibiting the effect at the peak density weighted by sample size. Clusters were reported using *Mango*.

The meta-analysis for negative associations of ghrelin identified similar regions in the density effect map (including putamen, pallidum, substantia nigra, caudate, thalamus, parahippocampal gyrus, and amygdala), yet no region survived extent-corrected threshold (*p* < .01). However, this analysis only included coordinates from 7 different contrasts which is too low to make robust inferences about convergence. Therefore, we discuss differential findings for ghrelin in the systematic literature review.

#### 1.3. GLP-1 MKDA: effects of GLP-1 do not survive correction

To estimate the hypothesized reward-suppressing effect of GLP-1, we conducted a meta-analysis for negative associations of GLP-1 with reward-related brain responses. In contrast to ghrelin, no cluster survived correction for multiple comparisons which could be due to the lower number of eligible contrasts. Descriptively, there was an overlap of identified coordinates with the neuromodulatory effects of ghrelin (Figure 2, Panel c). Since only three contrasts reported a positive association, we did not test for a positive association of GLP-1 with reward-related brain responses.

### 2. Systematic literature review

#### 2.1. Ghrelin

Studies looking at changes in BOLD activation in reward-related areas in association with ghrelin mainly focused on a restricted range of populations. Of the 25 papers included for the ghrelin subsample, 12 studies reported data from healthy participants, 6 for individuals with overweight and obesity, two for participants with addiction, and one for patients with bariatric surgery, anorexia nervosa and Prader-Willi Syndrome each. Most papers investigated correlations of plasma ghrelin levels in the fasted, or fed state, and less papers investigated the change in ghrelin levels following a meal (e.g., Sun et al., 2014). Studies also differed with respect to ghrelin, with some reporting on either ghrelin (10), des-acyl ghrelin (2), total ghrelin (16), or a combination. In four studies, ghrelin (versus placebo) was injected intravenously (Farokhnia et al., 2018; Goldstone et al., 2014; Han et al., 2018a; Malik et al., 2008).

#### 2.1 The modulatory role of ghrelin on reward responses across different populations

##### Healthy individuals

As detailed in the Supplementary Material (S1), 10 studies with healthy participants were included in the meta-analysis since they reported contrasts with a positive association with ghrelin. These studies applied different designs, such as administering a caloric load (Kroemer et al., 2013; Sun et al., 2014), or inducing caloric restriction for a prolonged period of time (Jakobsdottir et al., 2016). One study asked participants to either distance themselves or indulge on the images presented during a food cue reactivity task (Janet et al., 2021). Furthermore, two studies in healthy participants also reported negative associations with ghrelin (Iven et al., 2019; Jakobsdottir et al., 2016). These contrasting findings might be explained by substantial differences in study design, sample, and reward phase. For example, one of the studies applied an 8-week intervention with caloric restriction, testing participants both before and after this intervention while fasted and satiated (Jakobsdottir et al., 2016). Data from the fasted condition are in line with findings from other studies, while only correlations in the satiated state show results in the opposite direction (Jakobsdottir et al., 2016). Moreover, they only tested postmenopausal women, which have been reported to show larger decreases in ghrelin immediately after food intake (Stojiljkovic-Drobnjak et al., 2019). Another study (women only) administered the bitter tastant quinine-hydrochloride intragastrically before performing a drink taste with rewarding chocolate milkshake (Iven et al., 2019). Whereas quinine-hydrochloride decreased ghrelin levels, it increased most consummatory reward responses in regions of the motivational system.

Finally, three studies injected ghrelin intravenously (Goldstone et al., 2014; Han et al., 2018a; Malik et al., 2008). Ghrelin administration in the fed state increased OFC and hippocampus activation similar to fasting (i.e., endogenous ghrelinemia) (Goldstone et al., 2014). In contrast to most studies that assessed anticipatory reward responses, Han and colleagues (2018a) used a reinforcement schedule to induce reward-prediction errors while assessing reward learning. This allowed them to model reward-prediction error related activity and expected value assigned to conditioned stimuli, instead of relating ghrelin to reward responses *per se*. In addition, they found that ghrelin strengthened hippocampus-ventral striatum coupling during food conditioning (Han et al., 2018a).

Notably, several studies also reported no significant correlation between ghrelin levels and reward responses (Belfort-DeAguiar et al., 2016; Goldstone et al., 2014; Page et al., 2013; Rihm et al., 2019; Wever et al., 2021). Rihm and colleagues (2019) assessed anticipatory reward responses in a value-based decision-making task. Whereas they found increased ghrelin levels after sleep deprivation, they were not associated with reward responses. One of these studies initially found significant positive correlation between fasting ghrelin levels and activity in the left inferior occipital gyrus and right superior occipital gyrus, but these results did not survive outlier correction (Wever et al., 2021). Whereas in one study the administration of ghrelin (versus placebo) in the fasted state increased anticipatory responses, they did not find a significant correlation between plasma ghrelin levels and reward responses at individual sessions (Goldstone et al., 2014). These findings constitute an important step towards a clearer definition of the role of ghrelin on reward-related activation, but also highlight the need for more studies targeting different phases of reward.

##### Overweight and obesity

Since ghrelin may contribute to food seeking and, perhaps, “overindulgence” during eating, it has received much attention concerning potential association with overweight and obesity. However, associations with reward responses in participants with overweight and obesity are less consistent. In one study, positive correlations were found between ghrelin levels and reward responses in the medial PFC, caudate and visual cortex (Neseliler et al., 2019). Two more papers focused on specific areas and reported no significant correlation with ghrelin levels (Sewaybricker et al., 2021; Steele et al., 2015). Sewaybricker and colleagues (2021) tested couples of twins in which at least one individual had obesity (75% of the sample is reported to have obesity). They focused on dorsolateral PFC activity and found no association with changes in ghrelin levels (Sewaybricker et al., 2021). Another study compared patients with obesity and hypothalamic damage, patients without obesity and hypothalamic damage, controls with obesity and normal-weight controls (Steele et al., 2015). To assess a possible correlation between reward responses and ghrelin levels, they limited their analysis to the insula and found no significant correlation in any of the groups (Steele et al., 2015). One study that assessed whole-brain activations did not find any association between fasting ghrelin levels and anticipatory food reward responses (Perakakis et al., 2021). Finally, two other studies found negative correlations with activation in dlPFC, right caudate, limbic and paralimbic regions (Bogdanov et al., 2020; Karra et al., 2013). However, one study used a guessing task involving monetary rewards instead of food rewards, and the final reward received by the participants did not depend on wins and losses during the task, thereby rendering the choices during the task hypothetical which might not resemble tasks that assess anticipatory or consummatory reward facets more directly (Bogdanov et al., 2020). In the second study, individuals homozygous for the fat mass and obesity associated gene (FTO) (rs9939609 A allele) were compared with individuals without the risk allele, showing that in participants with the “obesity risk” gene, anticipatory responses were positively correlated with fasting acyl-ghrelin levels and negatively correlated with satiated levels. In contrast, participants without the risk allele showed correlations in the opposite direction (Karra et al., 2013). Thus, in line with mixed findings on the potential of medication targeting ghrelin to induce changes in body weight (Gupta et al., 2021), the current body of evidence shows limited convergence for ghrelin increasing reward responses in obesity.

##### Bariatric surgery

Studies in participants with overweight and obesity have also investigated the effects of weight-loss surgeries such as Roux-en-Y gastric bypass (RYGB), bariatric surgery, massive weight loss, or sleeve gastrectomy. One study investigated loss of appetite following laparoscopic sleeve gastrectomy (LSG) through a food cue reactivity task (Li et al., 2019). They found a correlation between attenuated dorsolateral PFC activity to high-caloric versus low-caloric food and reduced ghrelin levels in LSG patients after surgery (Li et al., 2019).

##### Drug addiction

Addiction to drugs has been tightly linked with altered activity in reward-related areas of our brain (Farokhnia et al., 2018; Koob et al., 2016). In patients with alcohol dependence, visual alcohol cues were found to increase anticipatory reward responses in distributed brain clusters, including ventral striatum during their first 21 days of controlled abstinence (Koopmann et al., 2019). Crucially, these reward-induced responses correlated positively with acyl ghrelin and the ventral striatal response fully mediated the effect of acyl ghrelin on alcohol craving (Koopmann et al., 2019). Moreover, administering ghrelin intravenously in participants with alcohol-dependence increased the anticipatory alcohol signal in the amygdala during alcohol clamp, and modulated an anticipatory food signal in the medial orbitofrontal cortex and NAc differentially, depending on alcohol consumption (Farokhnia et al., 2018).

##### Anorexia nervosa

Perhaps surprisingly, only one eligible study investigated ghrelin during reward tasks in anorexia nervosa (Holsen et al., 2014). They tested (female) patients with anorexia nervosa, individuals who recovered from anorexia nervosa and healthy-weight controls using a food cue reactivity task. The weight-recovered group showed a negative correlation between fasting levels of acyl ghrelin and brain responses to images of high-caloric food (vs. objects) in the left hippocampus. No such correlation was found in the two other groups. In contrast, they found a positive association between fasting levels of ghrelin and food reward responses in the amygdala, hippocampus, anterior insula, and OFC in healthy control participants (Holsen et al., 2014).

##### Major depressive disorder

Individuals with obesity often also report concurrent depressive symptoms (Mannan et al., 2016). These two conditions are interconnected, making the study of their shared neurobiological mechanisms fundamental to provide more effective interventions (Milaneschi et al., 2019). One study on patients with major depressive disorder compared patients with hyperphagic versus hypophagic symptoms with healthy controls, finding differential effects of ghrelin depending on somatic symptom profile (Cerit et al., 2019). Specifically, participants with hyperphagic depression showed a positive association between postprandial change in ghrelin levels and anticipatory food reward responses in the VTA and left hypothalamus, while participants with hypophagic depression showed a negative association in the right hypothalamus (Cerit et al., 2019).

##### Prader-Willi syndrome

Prader-Willi syndrome is a genetic disorder characterized by a constant sense of hunger, leading to hyperphagia and often to comorbid obesity (van Nieuwpoort et al., 2021). Therefore, studying the role of metabolic hormones in hyperphagia related to this disorder might lead to crucial insights about derailed reward responses. One study assessed changes in anticipatory food reward responses in adults with Prader-Willi Syndrome versus healthy controls, but they found no correlation between these changes and levels of fasting ghrelin (van Nieuwpoort et al., 2021).

Overall, these studies in disorders that are characterized by motivational dysfunctions highlight the potential role of metabolic hormones (Cerit et al., 2019; Holsen et al., 2014; Karra et al., 2013), although more studies are necessary to potentially pave the way for new interventions.

#### 2.2. GLP-1

Studies looking at changes in reward responses in association with GLP-1 mainly focused on metabolic disorders. Of the 20 papers included for the GLP-1 subsample, three studies included data from healthy participants, 10 included individuals with overweight and obesity, seven included individuals with T2D, four studies investigated GLP-1 in patients with gastric bypass, one included twins, and, one study included individuals with acquired hypothalamic damage. Most studies investigated correlations with plasma GLP-1 levels or administered GLP-1 agonists (6) or GLP-1 antagonists (5), or a combination (1).

##### 2.2.1. The modulatory role of GLP-1 on reward responses across different populations

###### GLP-1 in obesity and T2D

Obesity is mechanistically linked with T2D since individuals with a diagnosis of T2D are often obese and it is the main risk factor for developing T2D (Leitner et al., 2017). GLP-1 and GLP-1R agonists have been shown to inhibit glucagon release and enhance glucose-dependent insulin secretion (Brunton & Wysham, 2020). To this end, associations of GLP-1 with anticipatory and consummatory food reward responses in patients with T2D and concurrent obesity have received much attention. GLP-1 agonists showed effects on reward responses during anticipation and consumption of food rewards in lean participants (caudate, OFC), in participants with obesity (OFC) and in participants with T2D (putamen, insula, amygdala) (van Bloemendaal et al., 2015a). Moreover, GLP-1 changes in healthy participants and individuals with obesity were negatively correlated with anticipatory responses to food reward in the OFC after glucose intake (Heni et al., 2015). Three studies investigated the effect of liraglutide, an approved GLP-1 agonist for the treatment of obesity. Liraglutide increased GLP-1 and gastric inhibitory peptide plasma levels, whereas total ghrelin plasma levels were not affected (Farr et al., 2016a; Farr et al., 2016b). Liraglutide administration did not lead to (short-term) changes in anticipatory food reward responses in the fed state (Farr et al., 2016b). However, in a fasted state, liraglutide decreased anticipatory food reward responses in the parietal cortex, insula, and putamen (Farr et al., 2016a). In addition, increased GIP levels correlated with reduced reward responses in the insula (Farr et al., 2016b). Baseline scores of emotional eating affected the early sensitivity to liraglutide treatment in participants with T2D, and emotional eating was associated with attenuated liraglutide-induced decreases in anticipatory food reward responses in the amygdala, insula, and caudate, as well as attenuated increases in consummatory responses in the insula and caudate (van Bloemendaal et al., 2015b; van Ruiten et al., 2022). However, another study found no significant correlation between endogenous GLP-1 levels and anticipatory activation for food cues (Perakakis et al., 2021). Interestingly, one study reported no correlation between GLP-1 and DLPFC response to food cues, but found that the interaction between GLP-1 levels and DLPFC response robustly predicted body weight change (Maurer et al., 2019). These findings confirm the fundamental role played by GLP-1 in the treatment of obesity and T2D, but suggest the need for further investigation of its exact mechanisms of action, as well as of possible differential effects of circulating levels of GLP-1 and administration of GLP-1R agonists.

To block the actions of endogenous GLP-1, GLP-1 antagonists such as exendin-9-39 have been used. Such antagonists have been shown to prevent meal-induced reductions of anticipatory food reward responses in bilateral insula, left putament and right OFC in patients with T2D (ten Kulve et al. 2015; van Bloemendaal et al., 2014), and in right insula, amygdala and left OFC in patients with obesity (van Bloemendaal et al., 2014). Similarly, in healthy individuals, food reward responses in the right insula were attenuated, but this change was not significant. Based on the results, ten Kulve and colleagues (2015) argue that endogenous GLP-1 is involved in satiating effects after meal intake in obese patients with T2D (ten Kulve et al. 2015).

###### GLP-1 and bariatric surgery

In both animal and human studies, GLP-1 levels were found to be higher after RYGB and vertical sleeve gastrectomy (Baboumian et al., 2019; Hutch & Sandoval, 2017). Moreover, anticipatory food reward responses after RYGB were blunted in the caudate as well as during consummatory responses in the insula. This was consistent with findings after GLP-1 antagonist administration, which resulted in larger activations indicating that GLP-1 antagonists may reduce modulatory effects of RYGB in participants with obesity (ten Kulve et al., 2017). Reduced anticipatory responses to high caloric food were also found in the inferior temporal and right middle occipital gyri, while enhanced anticipatory responses were found in the medial prefrontal gyrus (Baboumian et al., 2019). Similar tendencies were found by Goldstone and colleagues (2016), but correlations between GLP-1 levels and anticipatory food reward responses failed to reach significance (Goldstone et al., 2016). Relatedly, before RYGB, a GLP-1 antagonist led to an increase in basal network connectivity and a decrease in functional connectivity in the frontoparietal network in patients with obesity (van Duinkerken et al., 2021). RYGB also reduced functional connectivity in the right OFC. After surgery, the GLP-1 antagonist increased connectivity in the default mode network relative to pre surgery (van Duinkerken et al., 2021). These findings provide useful insights on how GLP-1 might influence metabolic changes after RYGB and could help defining a more targeted intervention to facilitate post-surgery improvements.

##### 2.1.2. GLP-1 across different reward facets

Since the convergence of meta-analytic results for GLP-1 was too low to provide support for a neuromodulatory role in food reward processing, it may be instructive to compare specific phases. Notably, the effects of a GLP-1 receptor agonist during anticipatory and consummatory phases of food reward revealed opposing changes in brain responses, reducing anticipatory while enhancing consummatory reward responses (van Bloemendaal et al., 2015a). Specifically, GLP-1 receptor activation decreased anticipatory food reward responses in the OFC, putamen, left insula, and left amygdala, while it increased consummatory reward responses in the right OFC, putamen, left insula, left amygdala, and right caudate (van Bloemendaal et al., 2015a). Similar decreases in anticipatory reward responses were found in response to glucose intake (Dorton et al., 2018; Heni et al., 2015) and accompanied by an overall reduction in subsequent ad libitum food intake (van Bloemendaal et al., 2015a). Importantly, a GLP-1 receptor antagonist blocked these response patterns, supporting the idea that these effects were indeed mediated by GLP-1 receptors (ten Kulve et al., 2017). Moreover, GLP-1 antagonists were also found to enhance resting state functional connectivity between the hypothalamus and the left lateral OFC, and the left amygdala (Meyer-Gerspach et al., 2018). Still, such a differential role of GLP-1 during anticipation vs. consummation is not supported by studies looking at longer term changes in GLP-1 after RYGB (ten Kulve et al. 2017). These reward phases are also linked since consummatory signals will depend on anticipated benefits (Kroemer & Small, 2016). Consequently, discerning specific patterns of activation pertaining to phases may provide mechanistic insights that may facilitate future applications across various disorders.

## Discussion

To tune reward-related behavior according to metabolic demands, gastrointestinal hormones communicate with the central nervous system as part of the gut–brain axis. Beyond their homeostatic functions, we show the modulatory role of the circulating gut-based peptides ghrelin and GLP-1 in enhancing and reducing reward responses, respectively. Our review is the first systematic literature search and meta-analysis to evaluate the published evidence on the modulatory role in humans as measured by fMRI. First, using an MKDA meta-analysis, we demonstrated that reward responses are elevated by ghrelin in crucial regions of the reward circuit whereas there is currently not enough convergence for studies on GLP-1. Second, we highlighted how ghrelin and GLP-1 differentially modulate reward phases, with a specific focus on different clinical populations including obesity, T2D, eating disorders, alcohol use disorder, and depression. Consequently, our systematic review and meta-analysis lends support to the hypothesized potential of gastrointestinal hormones to modulate reward responses in humans. Although preliminary studies across various metabolic and mental disorders flank these promising findings regarding potential treatments of motivational dysfunction, we also highlight the need to investigate reward phases beyond anticipation to fully capitalize on the potential of gut-based interventions, specifically for mental disorders.

Our meta-analysis showed spatial convergence of neuroimaging studies which reported a positive association with ghrelin in several regions: the NAc, globus pallidus and dorsal mid insula, thalamus, putamen, right parahippocampal gyrus, amygdala, hypothalamus, midbrain, substantia nigra, and VTA. These areas are critically involved in reward-related pathways, and play an important role in motivation, goal-directed behavior as well as processing of different types of reward (Berridge & Robinson, 1998; Iven et al., 2019; Münte et al., 2017; Otis et al., 2019). The spatial convergence map for ghrelin resembles the dopamine motive system (Volkow et al., 2017), which is equipped with several metabolic receptors including GHSR and communicates with other networks that are involved in food intake, such as projections from the hindbrain and hypothalamus that forward homeostatic feedback (Howick et al., 2017; van Zessen et al., 2012; Volkow et al., 2017). Whereas fMRI does not allow inferences on neurotransmission, these results are well in line with work in animals (Decarie-Spain & Kanoski, 2021; Perello & Dickson, 2015). Moreover, we found overlapping density of effects for ghrelin and GLP-1 in striatal and dorsal mid insular regions. However, no regions survived correction for multiple comparisons for GLP-1. This overlap must be interpreted cautiously before more studies focusing on GLP-1 are available since the number of eligible contrasts for the meta-analysis and extracted peak coordinates were lower. Collectively, these findings provide encouraging support for the potential of gut-based interventions to rapidly alter reward responses since most currently available treatments of reward dysfunctions act more slowly (Frazer & Benmansour, 2002).

### Diversity of reward phases

Although reward involves distinct psychological and neurobiological components encompassing motivation (“wanting”), hedonic impact (“liking”), and learning (Berridge & Robinson, 2003), most studies focus on reward anticipation using food cue reactivity tasks. In fact, just seven out of the 29 studies investigated the consummatory phase of reward (Bogdanov et al., 2020; Han et al., 2018a; Sun et al., 2014; ten Kulve et al., 2017; van Bloemendaal et al., 2015a), and only one reward learning (Han et al., 2018a). Although some studies investigated the relationship between ghrelin or GLP-1 and behavioral motivation (e.g., Willingness to pay, Rihm et al., 2019; progressive-ratio alcohol self-administration, Farokhnia et al., 2018), none investigated motivational reward phases during fMRI. Crucially, the direction of the effect of GLP-1 may depend on the reward facet, such that GLP-1 attenuates anticipatory reward responses while it enhances consummatory responses (van Bloemendaal et al., 2015a). Whereas it is tempting to explain diverging results by referring to different reward phases, there is not enough evidence on reward phase specificity of gastrointestinal hormones yet due to the focus on anticipation. Nonetheless, a recent review including rodent studies supports the notion that GLP-1 has differential effects before compared to after a meal (Decarie-Spain & Kanoski, 2021). Taken together, future studies investigating the interaction of homeostatic and reward signals will benefit from incorporating several reward phases in the study design and analysis.

### Diversity of reward types

Studies in humans focused mostly on food reward and only a limited number of studies identified through the systematic literature search used other reward types such as alcohol or money. With respect to ghrelin, Han and colleagues (2018) reported no generalization of the effects of ghrelin administrations to non-food related odors and Bogdanov and colleagues (2020) found a negative association between total ghrelin and dorsolateral PFC responses to monetary wins. However, studies investigating ghrelin’s association with alcohol-related cues supports reward modulation beyond food reward (Bogdanov et al., 2020). With respect to GLP-1, a recent review by Eren-Yazicioglu et al. (2021) showed that GLP-1 treatment significantly reduced addictive-like behavior for drugs such as cocaine, alcohol, and nicotine in animals. However, none of the studies identified in this systematic literature review used rewards other than food for GLP-1. Investigating the generalization of existing results across different reward domains is encouraged to better understand the translational power of these findings.

### Diversity of clinical population

We highlighted a handful of studies that assessed the role of ghrelin and GLP-1 on reward responses in samples with metabolic and mental disorders. Most papers focused on either healthy participants (for ghrelin) or participants with obesity or T2D (for GLP-1). In contrast, metabolic disorders and disorders related to dysfunctions in reward processing (i.e., major depressive disorder and substance use disorders) have received less attention to date. This creates a translational gap, specifically for conditions where current treatment fail to achieve rapid improvements in symptoms (e.g., anhedonia in major depressive disorder; Treadway & Zald, 2011). Hence, research on associations between metabolic hormones and activation of reward-related areas during the different phases of reward processing in these populations is needed to evaluate potential applications as treatments (van Bloemendaal et al., 2015a).

### Limitations of the approach

Our meta-analysis and systematic review has limitations that should be considered when interpreting our findings.

First, many studies in the systematic review reported data from small samples of 10-15 individuals, particularly in clinical populations. Whereas sample sizes for studies included in the review were larger (Ghrelin: M(N)=23, Med(N)=27; GLP-1: M(N)=37, Med(N)=16) and considered by weighting coordinates, the overall sample size is still low. For ghrelin, we included 18 contrasts from 14 different studies with a combined sample size of 353 individuals. For the GLP-1 meta-analysis, we included 12 contrasts from 7 different studies with a combined sample size of 258 individuals. Increasing the sample size of future studies will likely improve the consistency of findings across studies as well.

Second, the heterogeneity of included studies was high. Due to the limited number of studies, we decided to include various tasks and populations in the meta-analysis. Potential differences were then considered as part of the systematic review. Despite the heterogeneity in study designs, this meta-analysis provides supporting evidence for spatial consistency for the modulatory effects of ghrelin on reward responses.

Third, coordinate-based meta-analysis are susceptible to various biases (Manuello et al., 2022). Although publication bias should always be considered (Jennings & Van Horn, 2012), neuroimaging studies are likely more susceptible to confirmation bias due to high analytical flexibility (Müller et al., 2018). Relatedly, some studies reported correlations for ghrelin or GLP-1 and anticipatory food reward activity for both high-caloric, low-caloric and non-food items, whereas others only reported selected contrasts increasing the risk of reporting bias. The analytical flexibility extends to additional decisions in the research process (“researcher’s degrees of freedom”; (Wicherts et al., 2016)) and includes different statistical thresholds that determine peak coordinates as input to any coordinate-based meta-analysis. With MKDA, the spatial consistency is not directly related to how many peaks were reported near a voxel. Instead, it is related to how many contrast maps activated near a voxel (Wager et al., 2009).

Fourth, coordinate-based meta-analyses are insensitive to non-significant results. To reduce this bias, we report all eligible studies in the systematic literature review, but this only help qualitatively, not quantitatively. Fifth, coordinate-based meta-analyses only assess spatial convergence of whole-brain findings and small-volume corrections skew the tests against a null hypothesis of random spatial convergence (Müller et al., 2018). To date, most studies do not share whole-brain statistical images. If contrast maps were openly shared as routine part of the publication process (e.g., on neurovault.org), image-based meta-analyses could overcome several limitations and provide more robust synthesis of research findings.

### Future direction and relevance for gut-brain interventions

More broadly, our review on the modulatory role of ghrelin and GLP-1 in reward processing adds to the growing evidence on the potential of gut-based interventions for motivational and emotional dysfunctions across a range of mental and metabolic disorders (Cryan et al., 2019; Grill, 2020; Horn et al., 2022; Howick et al., 2017). For example, in major depressive disorder, anhedonia is a cardinal symptom that is difficult to treat with conventional treatments (Argyropoulos & Nutt, 2013). In addition, patients with major depressive disorder may suffer from either increases or decreases in appetite and weight during a depressive episode (Milaneschi et al., 2019), which is associated with differences in energy metabolism (Simmons et al., 2018) and function of the reward circuit (Kroemer et al., 2022; Simmons et al., 2016). Beyond research on circulating gut-derived peptides, there has been a surge of interest in methods targeting the link between energy metabolism and reward via vagal afferent projections (Maniscalco & Rinaman, 2018; Tellez et al., 2013). Illustratively, recent studies have shown that non-invasive vagus stimulation boosts motivation (Neuser et al., 2020), improves mood (Ferstl et al., 2020; Kraus et al., 2007), and increases coupling between gastric myoelectric activity of the stomach and brain signals (Müller et al., 2022), which is in line with findings on vagal signaling derived with invasive methods in rodents (Cao et al., 2022; Han et al., 2018b). Moreover, various interventions have shown to exert beneficial effects on behavior via alterations in the microbiome (Cryan et al., 2019). Notably, neuromodulatory effects elicited by the microbiome are largely dependent on vagal signaling (Bravo et al., 2011; Breit et al., 2018; Fulling et al., 2019). Despite the large unmet demand for rapidly acting interventions, more research on the interaction between these facets of gut-based modulation is necessary to fully capitalize on the considerable potential of gut-based interventions in psychiatry, specifically for motivational dysfunctions.

## Conclusion

Hunger and satiety are potent modulators of reward responses, but can we harness the neuromodulatory potential of metabolic signals provided by gastrointestinal hormones ghrelin and GLP-1 in humans as well? Our systematic review and meta-analysis shows that ghrelin enhances reward responses within the motivational circuit as hypothesized. Although the convergence across studies was still too low for GLP-1, we identified an overlap of reported activations with the effects of ghrelin as well as potential phases-dependent effects that warrant further emphasis in future studies. Collectively, the current set of published evidence provides support for the hypothesized neuromodulatory potential of gastrointestinal hormones in altering reward responses. Still, to pave the way for more widespread clinical applications of gut-based interventions, future research will need to shift emphasis on more reward phases and clinical populations, ideally using larger samples with publication of the associations regardless of their threshold as full maps and regardless of a potential confirmation or rejection of the hypothesized action on reward responses.

## Supporting information

S1_Overview_of_included_papers.pdf

S2_Coordinates_for_meta-analysis.xlsx

S3_Additional_Results_Ghrelin_meta-analysis.docx

## Data availability statement

All papers included in the meta-analysis and systematic review are listed with additional information in Supplementary Material S1, and online with options to filter the database (https://tinyurl.com/gut-brain-reward). Data used for the meta-analysis was derived from papers cited in the reference section and contacting respective authors. To facilitate future meta-analyses, we provide all extracted peak coordinates along with author, publication date, and sample size in Supplementary Material S2. In addition to the results reported here, we provide the density effect map and thresholded maps for the ghrelin meta-analysis at https://identifiers.org/neurovault.collection:13247.

## Acknowledgement

The work was supported by the Deutsche Forschungsgemeinschaft (DFG) grants KR 4555/7-1 and KR 4555/9-1.

## Author contributions

NBK was responsible for the study concept and design. CV conducted the systematic literature search. CV & CS synthesized the literature. CS performed the meta-analysis and CV, & NBK contributed to analyses and their interpretation. CS & CV wrote the initial draft. All authors contributed to the interpretation of findings, provided critical revision of the manuscript for important intellectual content, and approved the final version for publication.

