## Supplementary material for "How gut hormones shape reward: a systematic review of the role of ghrelin and GLP-1 in human fMRI": S1_Overview_of_included_papers.pdf

Database of included stu...

Filter Sort

Author & Year | Hormonal assessment or ... | Reward Phase | Population | Reward Type | Inclusion Metaanalysis | Tags | + Add filter

| Author & Year | Title | Inclusion M... | Inclusion Sys... | Tags | Hormonal assessmen... | Reward Phase | Reward Type | Population |
|---|---|---|---|---|---|---|---|---|
| Baboumian et al., 2019 | Functional Magnetic Resonance Imaging (fMRI) of Neural Responses to Visual and Auditory Food Stimuli Pre and Post Roux-en-Y Gastric Bypass (RYGB) and Sleeve Gastrectomy (SG) | Yes | Include | GLP-1 | plasma GLP-1 | anticipation | Food | Roux-en-Y Gastric bypass, SG patients |
| Belfort-DeAguiar et al. 2016 | Food image-induced brain activation is not diminished by insulin infusion | No | Include | Ghrelin | plasma total ghrelin | anticipation | Food | Healthy |
| Bogdanov et al., 2020 | Reward-related brain activity and behavior are associated with peripheral ghrelin levels in obesity | No | Include | Ghrelin | plasma total ghrelin | consummatory | Money | Obesity |
| Cerit et al., 2019 | Divergent associations between ghrelin and neural responsivity to palatable food in hyperphagic and hypophagic depression | Yes | Include | Ghrelin | plasma acyl-ghrelin | anticipation | Food | Healthy, Depression |
| Dorton et al., 2018 | Influences of Dietary Added Sugar Consumption on Striatal Food-Cue Reactivity and Postprandial GLP-1 Response | No | Include | GLP-1 | plasma GLP-1 | anticipation | Food | Healthy |
| Farokhnia et al., 2018 | Exogenous ghrelin administration increases alcohol self-administration and modulates brain functional activity in heavy-drinking alcohol-dependent individuals | No | Include | Ghrelin | injection acyl-ghrelin, plasma acyl-ghrelin | anticipation, consummatory, effort | Food, Alcohol | Alcohol addiction |
| Farr et al., 2016 | Short-term administration of the GLP-1 analog liraglutide decreases circulating leptin and increases GIP levels and these changes are associated with alterations in CNS responses to food cues: A randomized, placebo-controlled, crossover study | No | Include | GLP-1, Ghrelin | plasma total ghrelin, GLP-1R agonist | anticipation | Food | Diabetes |
| Farr et al., 2016 | GLP-1 receptors exist in the parietal cortex, hypothalamus and medulla of human brains and the GLP-1 analogue liraglutide alters brain activity related to highly desirable food cues in individuals with diabetes: a crossover, randomised, placebo-controlled trial | Yes | Include | GLP-1 | GLP-1R agonist | anticipation | Food | Diabetes |
| Goldstone et al., 2014 | Ghrelin mimics fasting to enhance human hedonic, orbitofrontal cortex, and hippocampal responses to food | No | Include | Ghrelin | injection acyl-ghrelin, plasma GLP-1, plasma acyl-ghrelin | anticipation | Food | Healthy |
| Goldstone et al., 2016 | Link between increased satiety gut hormones and reduced food reward after gastric bypass surgery for obesity | No | Include | GLP-1 | plasma GLP-1 | anticipation | Food | Roux-en-Y Gastric bypass |
| Han et al., 2018 | Ghrelin Enhances Food Odor Conditioning in Healthy Humans: An fMRI Study | Yes | Include | Ghrelin | injection acyl-ghrelin | learning | Food | Healthy |
| Heni et al., 2015 | Dissociation of GLP-1 and insulin association with food processing in the brain: GLP-1 sensitivity despite insulin resistance in obese humans | Yes | Include | GLP-1 | plasma GLP-1 | anticipation | Food | Healthy, Obesity |

| Author & Year | Title | Inclusion M... | Inclusion Sys... | Tags | Hormonal assessmen... | Reward Phase | Reward Type | Population |
|---|---|---|---|---|---|---|---|---|
| Holsen et al., 2014 | Abnormal relationships between the neural response to high- and low-calorie foods and endogenous acylated ghrelin in women with active and weight-recovered anorexia nervosa | Yes | Include | Ghrelin | plasma acyl-ghrelin | anticipation | Food | Anorexia |
| Iven et al., 2019 | Intragastric quinine administration decreases hedonic eating in healthy women through peptidemediated gut-brain signaling mechanisms | Yes | Include | Ghrelin | plasma acyl-ghrelin<br>plasma total ghrelin<br>other substance administra | consummatory | Food | Healthy |
| Jakobsdottir et al., 2016 | Acute and short-term effects of caloric restriction on metabolic profile and brain activation in obese, postmenopausal women | Yes | Include | Ghrelin | plasma total ghrelin | anticipation | Food | Healthy |
| Janet et al., 2016 | Cognitive and hormonal regulation of appetite for food presented in the olfactory and visual modalities | Yes | Include | Ghrelin | plasma total ghrelin | willingness to pay<br>anticipation | Food | Healthy |
| Karra et al., 2013 | A link between FTO, ghrelin, and impaired brain food-cue responsivity | Yes | Include | Ghrelin | plasma acyl-ghrelin<br>plasma total ghrelin | anticipation | Food | Healthy |
| Koopmann et al., 2019 | Ghrelin modulates mesolimbic reactivity to alcohol cues in alcohol-addict[illegible] subjects: a functional imaging study | Yes | Include | Ghrelin | plasma acyl-ghrelin<br>plasma total ghrelin | anticipation | Alcohol | Alcohol addiction |
| Kroemer et al., 2013 | Fasting levels of ghrelin covary with the brain response to food pictures | Yes | Include | Ghrelin | plasma des-acyl ghrelin | anticipation | Food | Healthy |
| Li et al., 2019 | Reduced plasma ghrelin concentrations are associated with decreased brain reactivity to food cues after laparoscopic sleeve gastrectomy | Yes | Include | Ghrelin | plasma total ghrelin | anticipation | Food | bariatric surgery<br>Obesity |
| Malik et al., 2008 | Ghrelin modulates brain activity in areas that control appetitive behavior | Yes | Include | Ghrelin | injection ghrelin (no informa | anticipation | Food | Healthy |
| Maurer et al., 2019 | Interaction of circulating GLP-1 and the response of the dorsolateral prefrontal cortex to food-cues predicts body weight development | Yes | Include | GLP-1 | plasma GLP-1 | anticipation | Food | Obesity |
| Meyer-Gerspach et al., 2018 | Endogenous GLP-1 alters postprandial functional connectivity between homeostatic and reward-related brain regions involved in regulation of appetite in healthy lean males: A pilotstudy | No | | GLP-1 | plasma GLP-1<br>GLP-1R antagonist | resting state | | Healthy |
| Neseliler et al., 2019 | Neurocognitive and Hormonal Correlates of Voluntary Weight Loss in Humans | Yes | Include | Ghrelin | plasma acyl-ghrelin<br>plasma total ghrelin | anticipation | Food | Obesity |
| Nieuwpoort et al., 2021 | Food-Related Brain Activation Measured by fMRI in Adults with Prader–Willi Syndrome | No | Include | Ghrelin | plasma total ghrelin | anticipation | Food | Prader-Willi Syndrome |
| Page et al., 2013 | Effects of fructose vs glucose on regional Cerebral Blood Flow in Brain Regions Involved With Appetite and Reward Pathways | No | Include | Ghrelin<br>GLP-1 | plasma total ghrelin<br>plasma GLP-1<br>other substance administra | resting state | | Healthy |
| Perakakis et al., 2021 | Fasting oxyntomodulin, glicentin, and gastric inhibitory polypeptide levels are associated with activation of reward- and attention-related brain centres in response to visual food cues in adults with obesity: A cross-sectional functional MRI study | No | Include | GLP-1<br>Ghrelin | plasma GLP-1<br>plasma ghrelin (no informat | anticipation | Food | Obesity |

| | | | | | | | | |
|---|---|---|---|---|---|---|---|---|
| Rihm et al., 2019 | Sleep Deprivation Selectively Upregulates an Amygdala–Hypothalamic Circuit Involved in Food Reward | No | Include | Ghrelin | plasma total ghrelin, plasma acyl-ghrelin, plasma des-acyl ghrelin | anticipation | Food | Healthy |
| Sewaybricker et al., 2021 | Reassessing relationships between appetite and adiposity in people at risk of obesity: A twin study using fMRI | No | Include | GLP-1, Ghrelin | plasma GLP-1, plasma total ghrelin | anticipation | Food | Obesity, twins |
| Steele et al., 2015 | Cerebral activations during viewing of food stimuli in adult patients with acquired structural hypothalamic damage: a functional neuroimaging study | No | Include | GLP-1, Ghrelin | plasma acyl-ghrelin, plasma GLP-1 | anticipation | Food | acquired hypothalamic dama, Obesity |
| Sun et al., 2014 | The neural signature of satiation is associated with ghrelin response and triglyceride metabolism | Yes | Include | Ghrelin | plasma total ghrelin | consummatory | Food | Healthy |
| ten Kulve et al., 2015 | Endogenous GLP-1 mediates postprandial reductions in activation in central reward and satiety areas in patients with type 2 diabetes | Yes | Include | GLP-1 | GLP-1R antagonist | anticipation | Food | Diabetes, Obesity |
| ten Kulve et al., 2017 | Elevated postoperative endogenous GLP-1 levels mediate effects of Roux-en-Y gastric bypass on neural responsivity to food cues | Yes | Include | GLP-1 | plasma GLP-1, GLP-1R antagonist | anticipation | Food | Roux-en-Y Gastric bypass |
| van Bloemendaal et al., 2014 | GLP-1 Receptor Activation Modulates Appetite- and Reward-Related Brain Areas in Humans | Yes | Include | GLP-1 | GLP-1R agonist | anticipation | Food | Diabetes, Obesity, Healthy |
| van Bloemendaal et al., 2015 | Brain reward-system activation in response to anticipation and consumption of palatable food is altered by glucagon-like peptide-1 receptor activation in humans | No | Include | GLP-1 | GLP-1R agonist, GLP-1R antagonist | anticipation, consummatory | Food | Diabetes, Obesity, Healthy |
| van Bloemendaar et al., 2015 | Emotional Eating is Associated with Increased Brain Responses to Food-Cues and Reduced Sensitivity to GLP-1 Receptor Activation | Results reported | Include | GLP-1 | GLP-1R agonist | anticipation, consummatory | Food | Obesity, Diabetes |
| van Duinkerken et al., 2021 | Cerebral effects of glucagon-like peptide-1 receptor blockade before and after Roux-en-Y gastric bypass surgery in obese women: A proof-of-concept resting-state functional MRI study | No | Include | GLP-1 | GLP-1R antagonist | resting state | | Roux-en-Y Gastric bypass |
| van Ruiten et al., 2022 | Eating behavior modulates the sensitivity to the central effects of GLP-1 receptor agonist treatment: a secondary analysis of a randomized trial | Yes | Include | GLP-1 | GLP-1R agonist | anticipation | Food | Diabetes |
| Wever et al., 2021 | Associations between ghrelin and leptin and neural food cue reactivity in a fasted and sated state | Yes | Include | Ghrelin | plasma total ghrelin | anticipation | Food | Healthy, Obesity |

COUNT 39
