## Supplementary material for "How gut hormones shape reward: a systematic review of the role of ghrelin and GLP-1 in human fMRI": S3_Additional_Results_Ghrelin_meta-analysis.docx

Supplementary S3

| **Table 1.** Results of the multi-level kernel density meta-analysis reporting a positive association of ghrelin and BOLD activation using the height- and extent-based corrected threshold. | | | | | | | |
| --- | --- | --- | --- | --- | --- | --- | --- |
| Cluster | Region | x | y | z | Voxel | Volume mm^3^ | Maximum *P* |
| 1 | right amygdala, right parahippocampus | 22 | -4 | -22 | 2 | 16 | 0.30 |
| 2 | left pallidum | -14 | -4 | -8 | 202 | 1616 | 0.38 |
|  | substantia nigra | -12 | -16 | -12 | 32 |  |  |
|  | Left pallidum, medial globus pallidus | -16 | -2 | -6 | 170 |  |  |
| 3 | right thalamus | 4 | -16 | -2 | 40 | 320 | 0.35 |
| 4 | Left thalamus | -2 | -12 | 8 | 1 | 8 | 0.31 |
| 5 | right short posterior insular gyrus | 36 | 0 | 8 | 2 | 16 | 0.34 |
| 6 | Left putamen | -22 | 2 | -6 | 61 | 488 | 0.18 |
| Clusters that surpass the height-corrected threshold (p < .05) are reported (Cluster 1-5) along with subclusters. In addition, the clusters that pass the medium extent-based threshold (p < .01) that are not within 10 mm of the clusters for the height-corrected are reported (Cluster 6). Coordinates are Montreal Neurological Institute standard stereotaxic spaces. *Voxel* indicates the number of 2×2×2-mm voxels. *Maximum P* is the test statistic, indicating the maximum proportion of studies exhibiting the effect at the peak density weighted by sample size. | | | | | | | |
